# Implementation of PAR-CLIP to characterize RNA-protein interactions in prokaryotes at nucleotide resolution

**DOI:** 10.1101/2020.06.29.178624

**Authors:** Sandeep Ojha, Chaitanya Jain

## Abstract

The identification of RNAs that are recognized by RNA-binding proteins (RNA-BPs) using techniques such as “Crosslinking and Immunoprecipitation” (CLIP) has revolutionized the genome-wide discovery of RNA-BP RNA targets. Among the different versions of CLIP that have been developed, the use of photoactivable nucleoside analogs incorporated into cellular RNA has resulted in high efficiency photoactivable ribonucleoside-enhanced CLIP (PAR-CLIP). Nonetheless, PAR-CLIP has not yet been applied in prokaryotes. To determine if PAR-CLIP can be used in prokaryotes, we determined suitable conditions for the incorporation of 4-thiouridine (4SU), a photoactivable nucleoside, into *E. coli* RNA, and for the isolation of crosslinked RNA. Applying this technique to Hfq, a well-characterized regulator of small RNA (sRNA)-messenger RNA (mRNA) interactions, we showed that PAR-CLIP identified most of the known sRNA targets of Hfq, as well as functionally relevant sites of Hfq-mRNA interactions at nucleotide resolution. Based on our findings, PAR-CLIP represents an improved method to identify both the RNAs and the specific regulatory sites that are recognized by RNA-BPs in prokaryotes.

## Introduction

RNA-BPs constitute a large and diverse class of proteins that regulate nearly every aspect of RNA function inside the cell. Among the hundreds of such proteins, the specific functions of many are not well understood. An important goal towards defining the cellular roles of RNA-BPs is to identify the RNAs they interact with inside the cell – not only to better understand the biological functions of the RNA-BPs – but also the basis of RNA recognition through a systematic analysis of their RNA targets.

The first genome-wide method developed to identify the RNAs recognized by RNA-BPs was based on RNA immunoprecipitation (RIP) (1). In this method, the protein of interest, as well as its associated RNAs, are purified by using anti-RNA-BP antibodies. Thereafter, the RNAs are extracted and their identities are revealed either by sequencing or by hybridization to DNA microarrays. Over the past few years RIP has become largely supplanted by CLIP, a method that is conceptually similar to RIP, except that the RNA-BPs are first crosslinked to their RNA targets *in vivo* (2,3). The development of CLIP stemmed, in part, from concerns that the non-covalent nature of protein-RNA interactions in RIP could result in the loss of many bound RNAs, especially if extensive purification steps were employed.

Two different procedures for crosslinking have been applied for CLIP. In the first, a chemical crosslinker, such as formaldehyde, has been used (4,5). In the second approach, RNA-protein crosslinks are generated by irradiating cells with short-wavelength ultraviolet (UV) light (2,3). Despite widespread adoption, each method suffers from limitations that can reduce their effectiveness. With chemical crosslinking, a significant drawback is that the crosslinking reagents induce not only protein-RNA, but also protein-DNA and protein-protein crosslinks. Therefore, the RNAs identified can include those that are recognized by proteins that interact with the RNA-BP of interest. In contrast, the efficiency of crosslinking with standard UV-CLIP is low, which can impact the coverage of RNA targets identified using this method. Any improvements to these techniques, therefore, are expected to yield a more complete and accurate picture of RNA-BP targets inside the cell.

To improve on existing procedures, a new method, PAR-CLIP, was developed (6). In this method, cells are grown in the presence of a photoactivable nucleotide added to the growth medium, the most commonly used being 4SU. Incorporation of 4SU into RNA, followed by irradiation with long-wavelength UV light, has been reported to enhance crosslinking efficiency by 100-1000-fold relative to conventional CLIP (6). An additional benefit of PAR-CLIP is that crosslinked 4SU residues are misread as cytosine residues with high frequency during the reverse transcription step of sequence library preparation (6–8). Consequently, an analysis of T to C conversion frequency in the sequence data directly identifies the sites of cross-linking in the RNAs.

Although PAR-CLIP is now established in eukaryotes, to the best of our knowledge, it has not yet been implemented in prokaryotes. To determine the feasibility of this technique for prokaryotic organisms, we first identified suitable conditions for its use in *E. coli.* We then applied PAR-CLIP to Hfq, a well-characterized regulator of sRNA-mRNA interactions, and found that PAR-CLIP effectively identifies most of the sRNAs previously shown to be associated with Hfq. Through the use of RNA-Seq and quantitative proteomics, we further identified a number of mRNAs that were both regulated by Hfq and contained PAR-CLIP cross-linking sites. Significantly, we found that mutation of residues in the vicinity of the cross-linking sites reduced the extent of Hfq-mediated regulation in almost every case that was analyzed. We anticipate that the power of PAR-CLIP to identify biologically relevant cross-linking sites will be highly useful in the future to define the function of RNA-BPs inside the cell.

## Results

### Incorporation of 4SU into E. coli RNA

To determine whether 4SU can be incorporated into *E. coli* RNA, a wild-type strain (CJ2109) was grown in LB medium or in LB supplemented with 300 μM 4SU. RNA was isolated from mid-log cultures and digested with nucleases and phosphatases to nucleosides. The digested RNA was analyzed by using high performance liquid chromatography (HPLC) with readings at 260 nm (all nucleotides) or 330 nm (4SU). Both sets of RNAs gave similar peaks for the standard nucleosides at 260 nM (Fig. 1). However, with detection at 330 nm, the RNA samples isolated from cells grown with 4SU yielded a significantly higher signal compared to cells without 4SU. Based on peak heights and a comparison with defined nucleoside standards, we calculated that 1.3% of uridine residues in the latter RNA sample were replaced with 4SU. These experiments indicated that 4SU can be incorporated into RNA *in vivo*.

**Figure. 1.**
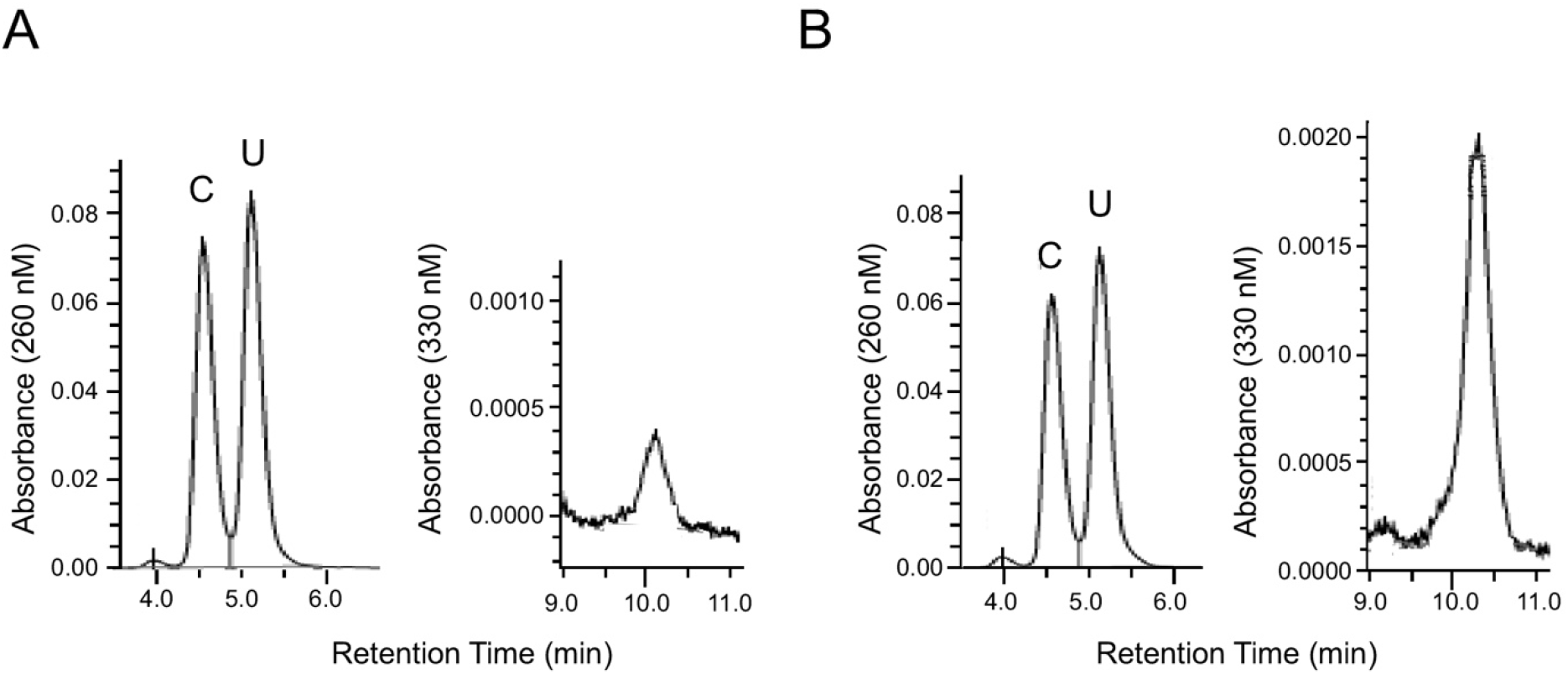
HPLC analysis. Total cellular RNA was isolated from cells grown in the absence (A) or presence of 300 μM 4SU (B), enzymatically digested to nucleosides, and the products were resolved by HPLC with detection at 260 nm or 330 nm. Only the regions corresponding to the retention times for uracil (U) and 4SU are shown. For the 260 nm trace, the mutually overlapping cytosine (C) peak is also included. A background 330 nm absorbance that is present in the RNA sample isolated from cells grown without 4SU is likely due to 4SU that is post-transcriptionally incorporated into tRNAs (28).

### PAR-CLIP on FLAG-tagged proteins

To determine whether 4SU containing RNA can be crosslinked to RNA-BPs, CJ2109 was transformed with pCJ1099, a plasmid that encodes the DEAD-box protein RhlE tagged with a 3X FLAG epitope at its C-terminal end. We have previously shown that RhlE binds to the 50S ribosomal subunit and to 70S ribosomes (9), and therefore, is likely an RNA-binding protein. The transformed strain (CJ2191) was grown to midlog phase in LB medium in the absence or the presence of different concentrations of 4SU in the medium, followed by harvesting of cells and irradiation with long-wavelength UV light (365 nm). The cells were lysed, incubated with beads that were coupled to anti-FLAG antibodies, and unbound proteins were removed through multiple washes. Thereafter, the crosslinked RNAs were fragmented on beads and labeled at the 5’ end with ^32^P. The labeled proteins were eluted from the beads, fractionated by using polyacrylamide gel electrophoresis (PAGE), transferred to nitrocellulose, and exposed to a phosphorimager screen to visualize the crosslinked RNAs.

A comparison of the signal derived from cells grown in the presence of different amounts of 4SU indicated that low levels of radiolabeled RNA were crosslinked to RhlE when cells were grown in the absence or the presence of 100 μM 4SU (Fig. 2A). Significant amounts of cross-linked signal was observed for cells grown with 250 μM 4SU and a further signal increase was observed with 500 μM 4SU. These results show that 4SU-containing RNAs can be crosslinked to proteins. Surprisingly, when cells were grown with 1 mM 4SU, the signal was found to be reduced. However, we noticed those cells exhibited slower growth, suggesting that the reduced efficiency of 4SU incorporation into RNA in this instance could be related to growth inhibition.

**Figure. 2.**
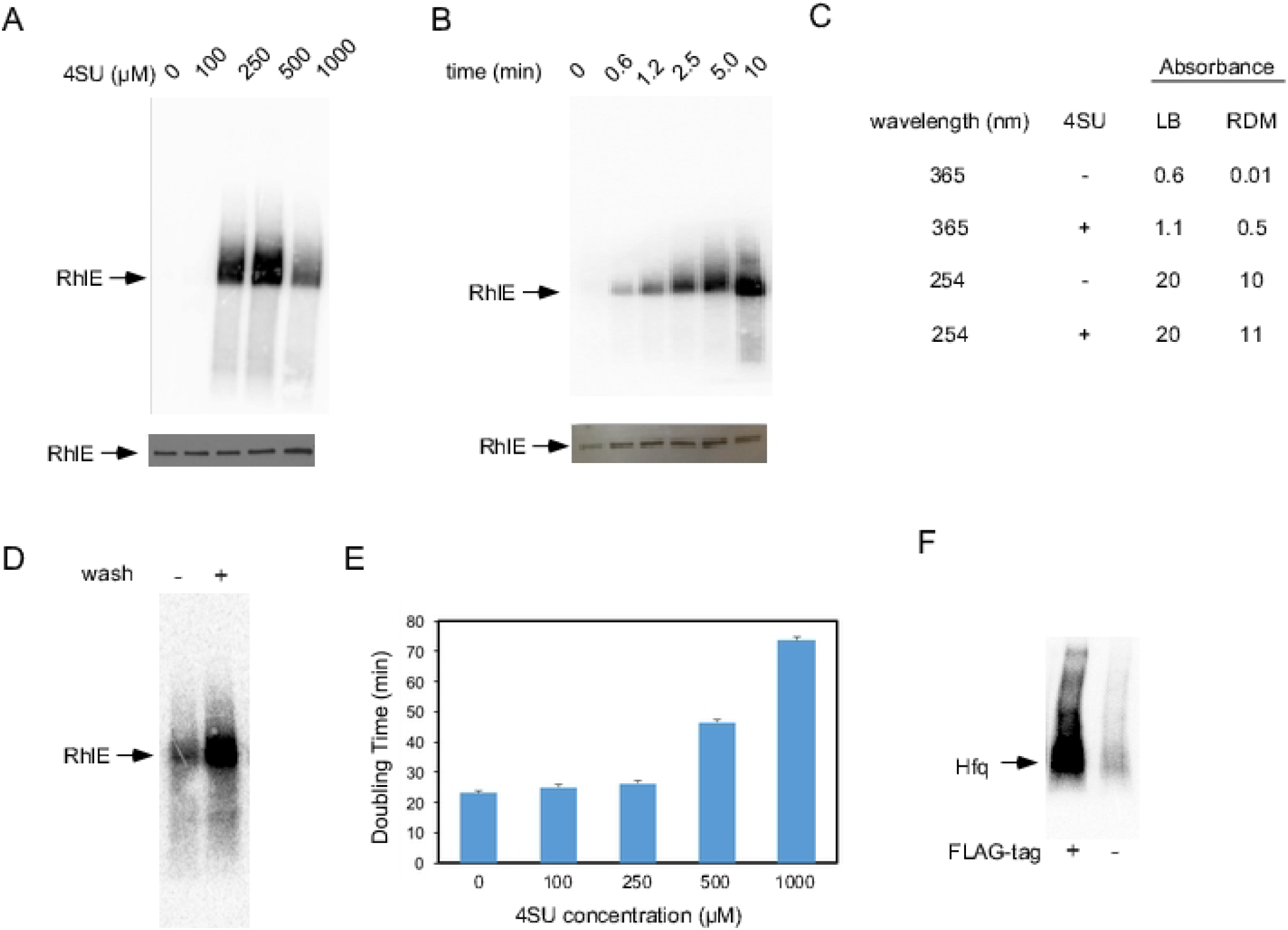
PAR-CLIP on FLAG-tagged proteins. (A) CJ2191 was grown in LB supplemented with different concentrations of 4SU and cells were crosslinked at 365 nm after harvesting. FLAG-tagged RhlE was purified by using anti-FLAG antibodies and the RNAs crosslinked to RhlE were radiolabeled. Proteins were resolved by gel electrophoresis and transferred to nitrocellulose filter. Top, the nitrocellulose filter was exposed to a phosphorimager screen to evaluate the extent of RNA crosslinking to RhlE; bottom, the filter was subsequently analyzed by Western blot using anti-FLAG antibodies. (B) The effect of UV irradiation time on cross-linking efficiency. CJ2191 was grown in LB medium and irradiated at 365 nm for different periods of time. The crosslinked RhlE was analyzed as in (A). Top, phosphorimager scan; bottom, western-blot analysis. (C) Optical density of LB medium and RDM. The absorbance of LB and RDM was measured at 365 and 254 nm in the absence or presence of 250 μM 4SU. (D) Effect of washing on cross-linking efficiency. CJ2191 was grown in LB medium containing 250 μM of 4SU and crosslinked prior to or after washing and resuspension in saline. (E) Effect of 4SU on cell growth. Strain CJ2109 was grown in RDM supplemented with the indicated concentrations of 4SU and cell doubling times were measured. The indicated standard errors are based on three replicates for each 4SU concentration. (F) PAR-CLIP on single-copy FLAG-tagged Hfq. Strains CJ2192 or CJ2109 were grown in RDM containing 250 μM of 4SU and were irradiated after harvest with 365 nm UV light for five minutes. FLAG-tag protein purification and RNA labeling was performed as in (A).

To optimize the crosslinking efficiency of 4SU-containing RNAs, cells were irradiated with 365 nm UV light for different amounts of time (Fig. 2B). Increasing the irradiation time for up to 10 minutes resulted in increased levels of crosslinking, indicating that the extent of crosslinking was not saturated even with the longest irradiation time used.

We were next interested in determining whether the efficacy UV irradiation could be limited by the medium used for growth. We compared the absorbance of LB medium, which is commonly used to grow *E. coli* cells, with a rich-defined medium (RDM). RDM, unlike LB, is a clear solution, and might be expected to absorb less long-wavelength UV light. We found that RDM was virtually transparent to 365 nm UV light, whereas LB medium displayed an absorbance of 0.6 (Fig. 2C), which corresponds to 75% absorbance of UV light by 1 cm of medium. As expected, the addition of 250 μM of 4SU increased absorbance in both growth media. Because the standard CLIP protocol uses 254 nm UV light, we also measured the absorbance of both LB and RDM at 254 nm. In each case, the absorbance was 10 or greater, likely because of the presence of nucleotides or their analogs in the media, which implies that when standard crosslinking is performed using 254 nm UV, the effective depth to which the UV light will penetrate is less than 1 mm. Although several factors determine the overall efficiency of cross-linking, the inability of 254 nm UV to penetrate the growth media might be considered a significant drawback of using such short wavelength light for cross-linking purposes.

To verify that absorbance by LB could be reducing cross-linking efficiency, CJ2191 was growth in LB/4SU and either cross-linked directly after cell harvest, or after pelleting the cells and resuspending them in saline (0.85% NaCl). Consistent with the results of Fig. 2C, we found that washing off the growth medium significantly increased the cross-linked signal (Fig. 2D). Nonetheless, given that many *E. coli* RNAs have short half-lives, and since a washing step would increase the time between harvesting cultures and cross-linking, we decided to adopt RDM as the growth medium of choice for our studies.

Because we had observed a reduction in cross-linking signal when cells were grown with 1000 μM of 4SU (Fig. 2A), we measured the effect of different concentrations of 4SU on cell growth. The addition of 4SU up to 250 μM to RDM had no significant effect on the growth rate, but increased 4SU concentrations was found to increase cell doubling time, indicating growth inhibition (Fig. 2E). In order to perform CLIP without perturbing growth, the downstream studies were performed using 250 μM of 4SU in the growth medium.

Finally, with an intention to generate CLIP data using a model RNA-BP that is expressed at physiological levels, we constructed a strain (CJ2192) in which the chromosomal locus expressing Hfq RNA-BP was modified to append a 3X FLAG-epitope at its C-terminus. Hfq is a well-studied regulator of gene expression, and multiple studies have shown that it binds to several regulatory sRNAs inside the cell and facilitates sRNA-mRNA interactions (10,11). CJ2192 was grown in the presence of 4SU, and the cells were crosslinked at 365 nm. To determine whether there is any non-specific background, the same procedure was also performed on the untagged strain, CJ2109. After crosslinking, the RNAs were analyzed by FLAG-tag protein purification and gel electrophoresis, as described above. With CJ2192, a strong signal consistent with the migration of Hfq could be observed (Fig. 2F). In contrast, very little signal was observed with the untagged strain, which could be due to free RNA or to other RNA-BPs that bind to the antibody-coupled beads (12). Based on these observations, we infer that sequence libraries made via PAR-CLIP on CJ2192 are expected to be highly enriched for RNAs that are specifically cross-linked by Hfq.

### Identification of Hfq-interacting sRNAs using PAR-CLIP

To convert the crosslinked RNA into sequencing libraries, strips containing the Hfq-crosslinked RNA were cut out from nitrocellulose filters and the RNA was released from the nitrocellulose strips using proteinase K to digest proteins. The purified RNAs were converted to sequencing libraries and sequenced on an Illumina HiSeq sequencing system. To assess any contribution from background RNA, PAR-CLIP was similarly performed on RNA derived from CJ2109 that was purified from the same region of the nitrocellulose filters as for FLAG-tagged Hfq. The resulting data were analyzed using the Galaxy bioinformatics platform (13).

We first assessed the fraction of reads that correspond to different classes of RNAs. A comparison of sequence reads between the two libraries revealed that the fraction of reads corresponding to tRNAs and rRNAs did not change appreciably between the two samples, but reads corresponding to non-coding RNAs (ncRNAs) were increased by over two-fold in the library from CJ2192 whereas mRNA reads were decreased (Fig. 3A). These changes are consistent the significant role of Hfq in interacting with sRNAs, which are included in the ncRNA category, and in targeting mRNAs for translational repression or degradation (14). Next, we assessed the frequency of T to C conversions, diagnostic of protein-4SU crosslinking, within the individual sRNAs. We found that 22 sRNAs showed significantly greater T>C conversion ratios in the library prepared from CJ2192, as compared to the control library (Fig. 3B). The majority of the sRNAs on this list have been identified as mRNA regulators (15), consistent with the role of Hfq as an effector of sRNA-mRNA interactions. To further define the effects of Hfq on these sRNAs, we performed RNA-seq on wild-type (wt) and Δ*hfq* strains (Supplementary Table 1). Among the sRNAs that were found to display enhanced T>C conversion ratios in the Hfq-tagged strain, we were able to quantify thirteen (Fig. 3C), half of which were found to be present at a lower level in the Δ*hfq* strain, which is consistent with Hfq being a stabilizing factor for many sRNAs (16).

**Figure 3.**
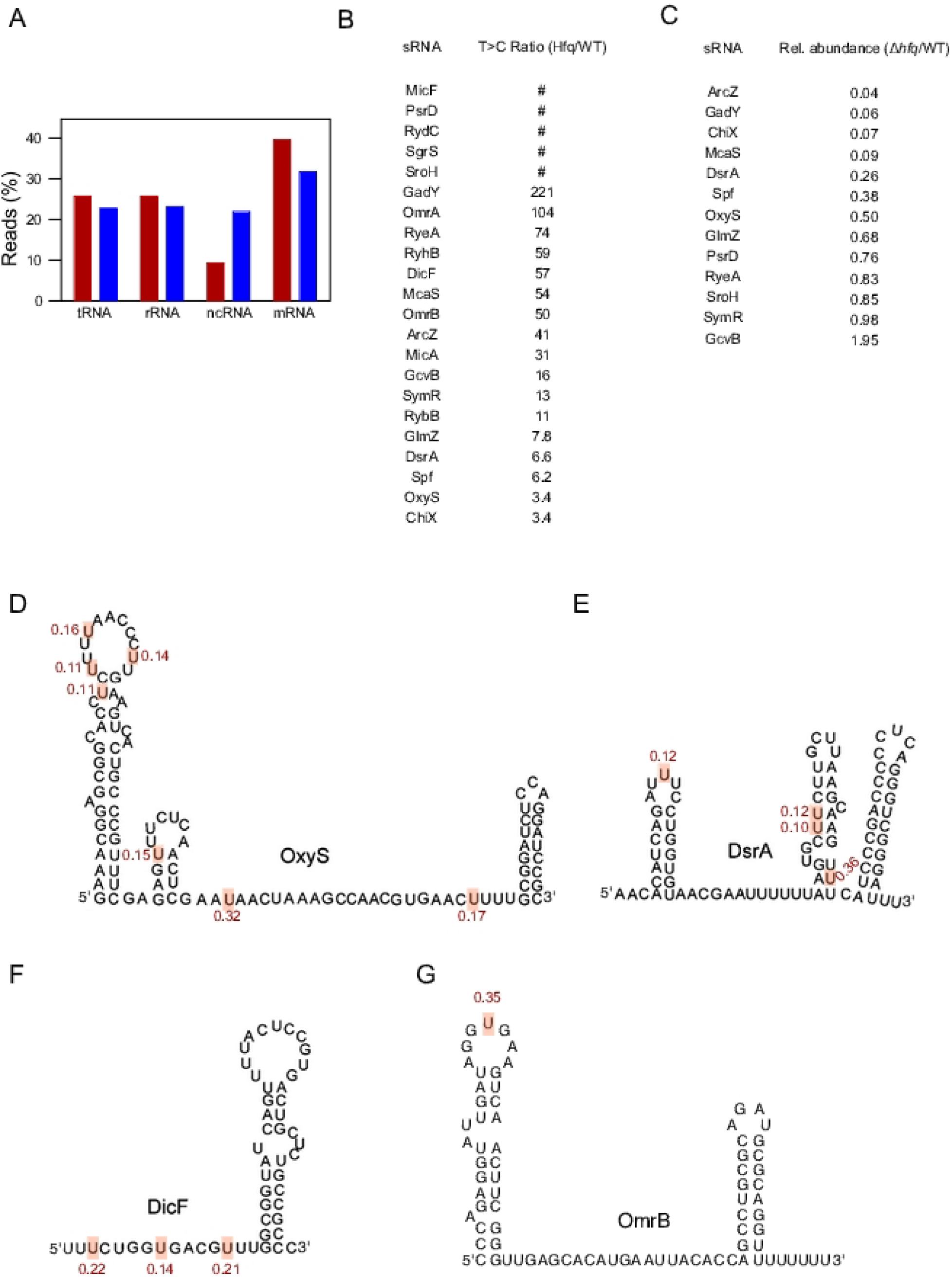
Analysis of sRNAs. (A) PAR-CLIP on Hfq enriches for ncRNAs. The percentage of uniquely mapping reads from PAR-CLIP libraries from strains CJ2109 (red bars) or CJ2192 (blue bars), which correspond to different categories of *E. coli* RNAs, are depicted. (B) sRNAs enriched in Hfq PAR-CLIP libraries. The sRNAs with cumulative T>C conversion frequencies that are two-fold or higher in PAR-CLIP libraries from strain CJ2192, relative to strain CJ2109, are listed. #, T>C changes were observed only in the CJ2192 library. (C) Effect of the Δ*hfq* mutation on sRNA abundance. The relative abundance of sRNAs in Δ*hfq* and wt strains, as determined by RNA-seq, is listed. (D-F) Sites of enhanced T>C conversions in the OxyS (D), DsrA (E), DicF (F) or OmrB (G) sRNAs. Only sites that displayed a two-fold or greater T>C conversion ratios in libraries made from strain CJ2192, as compared to strain CJ2109, are shown. The T>C conversion frequency at these positions is shown in red. The RNAs are depicted based on known structures or computationally derived structures using Mfold (29).

To define specific sites of interaction between Hfq and the sRNAs, we characterized T>C conversion ratios at the nucleotide level for each of the 22 sRNAs (Supplementary Table 2). We found the different sRNAs to contain between 1-11 positions that displayed enhanced T>C ratios, some examples of which are presented in Fig. 3D-G. For the OxyS sRNA, seven sites with increased T>C conversion ratios were identified, of which one, at position 68, showed a two to three-fold higher conversion ratio compared to the other positions (Fig. 3D). Consistent with the increased T>C conversion frequency observed at this position, it has been shown that Hfq binding to OxyS is mediated by residues near nt 70 (17). A second sRNA, DsrA, exhibited four positions with enhanced T>C conversion ratios (Fig. 3E), of which one, present at the base of the second stem-loop motif in DsrA, exhibited a three-fold higher ratio than the other sites, which suggests that Hfq might be binding to DsrA primarily through recognition of this stem-loop structure. Of significance, none of the six contiguous U-residues preceding this stem-loop exhibited T>C conversion changes, which is surprising since stretches of U residues have been identified as an Hfq-binding motif in several other RNAs (11). However, a lack of T>C conversions at these positions is fully consistent with a previous study that showed that mutation of this U-rich stretch has little effect on the interactions between Hfq and DsrA (18). A third sRNA, DicF, was found to exhibit increased T>C conversion frequencies only within a single-stranded region, whereas in the final example, a single position within the loop region of OmrB was identified (Figs. 3F and G). Based on the different contexts in which these sites are located on the sRNAs, it indicates that Hfq can bind to RNA within a variety of different RNA motifs.

### Identification of Hfq-regulated proteins

The RNA-Seq analysis also identified a number of mRNAs that were differentially expressed in Δ*hfq* strains (Supplementary Table 1). However, mRNAs that are translationally regulated by Hfq would not be expected to be identified in this population. To identify additional Hfq-regulated genes, we performed quantitative mass spectrometry analysis on cell extracts derived from wt and Δ*hfq* strains. With a view to identify Hfq-regulated genes under more than one condition, mass spectrometry was performed both on cells grown to mid-log phase and on cells growth to stationary phase (Supplementary Table 3), as the latter condition is known to increase the extent of regulatory control by Hfq through an increase in the concentration of Hfq and that of several sRNAs via the stabilization role of Hfq (19).

For cells grown to midlog phase, these analyses yielded a quantitation of 1561 proteins. Among these proteins, 35 showed a two-fold or greater increase in the Δ*hfq* strain, as compared to the wt strain, and 30 showed a two-fold or greater decrease (Fig. 4A). Similarly, for cells grown to saturated phase, 1641 proteins were quantified of which 45 were upregulated by two-fold or more in the Δ*hfq* strain, and 75 were downregulated by two-fold or more (Fig. 4B). A comparison of proteins that were upregulated under the two growth conditions a yielded significant overlap (Fig. 4C), but however, more than half of the proteins that were upregulated by two-fold or greater in stationary phase (24/45) were not upregulated to the same extent in mid-log phase. In a similar manner, the majority of proteins downregulated by two-fold or more in midlog phase were also downregulated during stationary phase, but the reverse was not true (Fig. 4D). These data suggest that the effects of Hfq on gene regulation are generally more significant during stationary phase. Consistent with this notion, among the proteins that were up- or down-regulated in both the midlog and stationary phase, a majority (28/42) showed a greater fold-effect in stationary phase (Supplementary Table 3). Nonetheless, we note that several proteins were significantly regulated in the midlog phase, but not in stationary phase.

**Figure 4.**
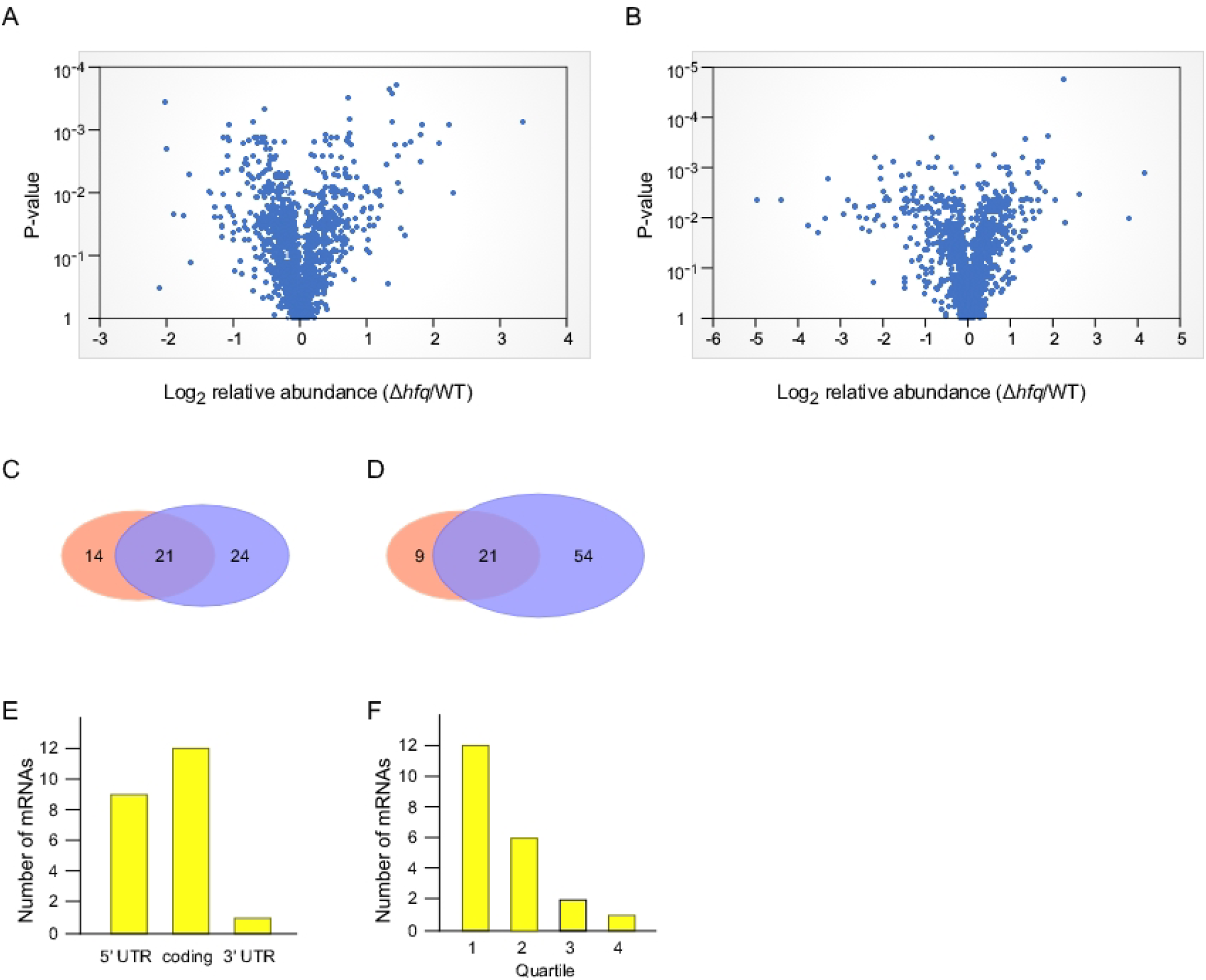
Analysis of hfq-regulated proteins. (A, B), Volcano plot of fold change in protein levels in the Δ*hfq* strain vs. P-value for cells grown in RDM to midlog phase (A) or stationary phase (B). P values are based on three replicates each. (C) Overlap of proteins upregulated in the Δ*hfq* strain upon growth to midlog phase (red ovals) or stationary phase (blue ovals). (D) Overlap as in (C) for proteins downregulated in the Δ*hfq* strain. (E) Location of sites with enhanced T>C conversion ratios for mRNAs corresponding to the proteins differentially expressed in the Δ*hfq* strain. In one case, (Dps) enhanced T>C conversion sites were identified both in the coding region and in the 3' UTR of its mRNA, and is included in both categories. (F) Abundance of mRNAs displaying enhanced T>C conversion ratios. mRNAs associated with enhanced T>C conversion ratios were assigned to one of four quartiles based on RNA-Seq data from a wild-type strain (Supplementary Table 1). Quartile 1 represents the 25% most abundant mRNAs, and quartile 4, the RNAs that are least abundant.

We next investigated whether the regulated proteins were associated with increase T>C conversion ratios within their mRNAs. Among the 143 proteins that were up-or down-regulated in either growth condition, 21 could be linked to increased T>C conversion ratios within their mRNA 5’ UTR, coding region and/or the 3’ UTRs (Fig. 4E, Supplementary Table 4). An incomplete match between the number of regulated proteins and those exhibiting increased T>C conversion frequencies within their mRNAs could be due either to some genes being regulated by Hfq indirectly or because some of the regulated mRNAs are present at a low abundance. We note that the PAR-CLIP libraries were prepared from cells in the midlog phase, whereas most of the proteins that were regulated were identified in stationary phase. To test the latter hypothesis, the 21 mRNAs with increased T>C conversion ratios were ranked based on their abundance in the RNA-Seq library. We found that the majority of RNAs were expressed at relatively high levels (Fig. 4F). Therefore, we expect that a significant fraction of the genes that are regulated by Hfq but could not be associated with T>C conversions might yet be direct targets whose low expression precluded the detection of T>C events.

### Validation of direct interactions between Hfq and Hfq-regulated RNAs

With the identification of several cross-linked mRNAs whose products were also differentially expressed in the Δ*hfq* strain, a key question that arose was whether the interaction sites identified by cross-linking accurately define the basis for regulation by Hfq. To address that question, we constructed *lacZ* reporter fusions with several of the regulated genes to test whether mutations at the cross-linking sites could affect regulation by Hfq. We found that some of the regulated genes, when fused to *lacZ*, were not appreciably regulated by Hfq in this context, which might be because some regions important for regulation were absent in these fusions or because the presence of *lacZ* sequences interferes with Hfq-mediated regulation. Nonetheless, among the fusions that were regulated to a significant degree, several were further analyzed, of which data for three are shown in Figs. 5A-5C, whereas data for additional constructs is summarized in Fig. 5D.

**Figure 5.**
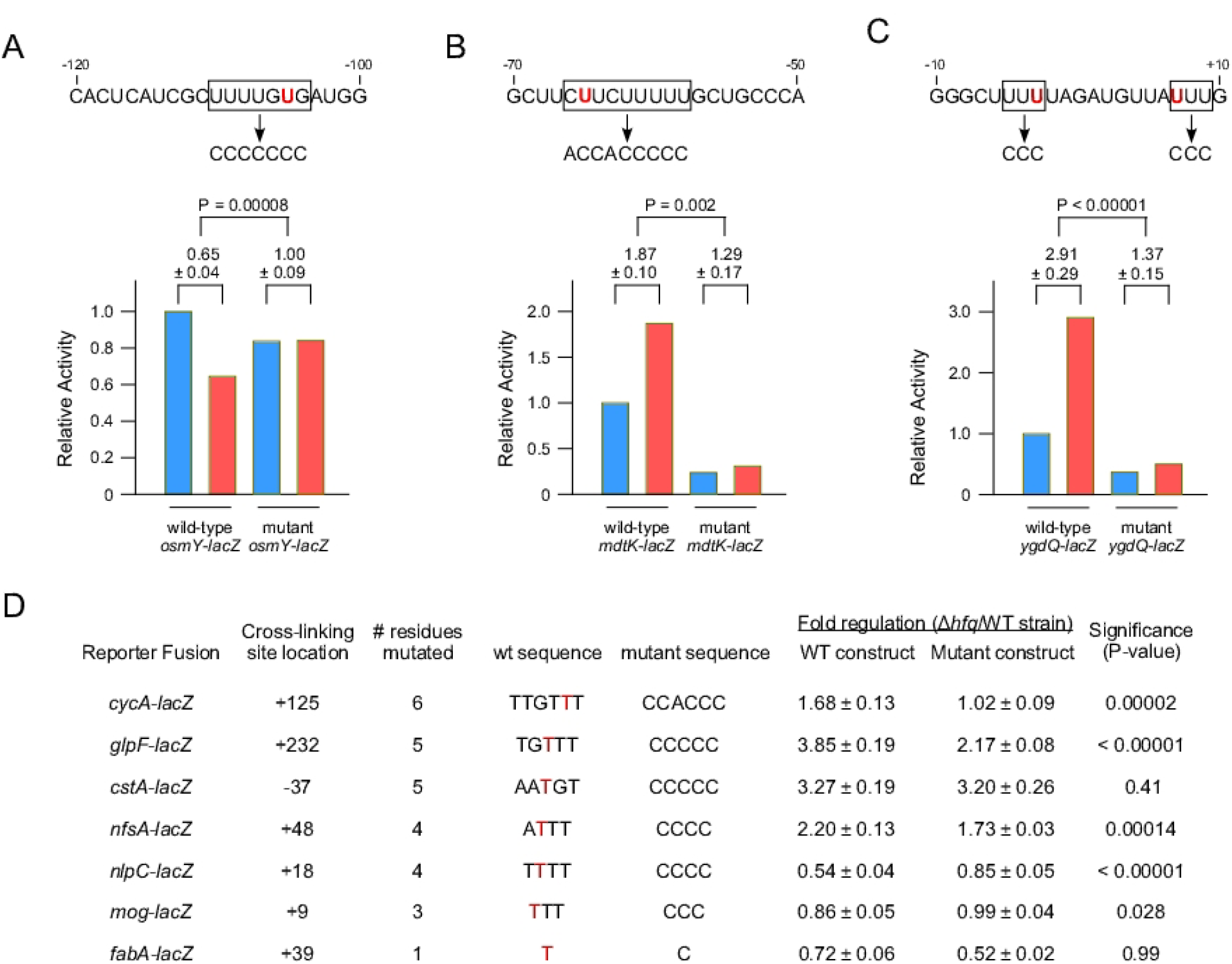
Mutation of residues near mRNA cross-linking sites reduces regulation by Hfq. (A-C) Top, sequences of the Hfq-regulated mRNAs for the *osmy* mRNA (A), *mdtK* mRNA (B) and *ygdQ* mRNAs (C). The cross-linked residues are shown in red and the sequences that were mutated depicted by arrows. Bottom, activity of the respective wild-type or mutant *lacZ* fusions. The *lacZ* activity of the fusions in wt and Δ*hfq* strains is depicted by blue and red bars, respectively. The values above the bars indicate the extent to which the *lacZ* activity of the fusions is regulated by Hfq. The P-values shown are based on the null hypothesis that mutations in the corresponding construct do not decrease the extent of Hfq-mediated regulation. (D) Analysis of additional Hfq targets identified by PAR-CLIP. The location of the cross-linking site relative to the start of the coding region and the number of residues mutated are indicated. The sequences of the residues near the cross-lining site in the wt and mutant constructs are shown, with the Hfq cross-linked residues in red. The extent of regulation of wt and mutant constructs, and P-values, were calculated as for sections A-C.

A first fusion that was analyzed was with *osmY*, which was found to be downregulated at the protein level in the Δ*hfq* strain and for which a single PAR-CLIP cross-linking site was identified in the *osmY* mRNA 5’ UTR, 106 nts upstream of the initiation codon. We noticed that the cross-linked site was included within a U-rich sequence comprised of five uridine residues within a seven nt tract (Fig. 5A), which could serve as additional sites of Hfq-RNA interactions. To determine whether those residues are responsible for regulation by Hfq, we made an *osmY-lacZ* mutant construct that converts each of these seven residues to a C residue. Strikingly, the activity of the wt *osmY-lacZ* fusion was found to be reduced by 1.5-fold in the Δ*hfq* strain, whereas the effect of Hfq on the activity of the mutant construct was completely abolished. On the basis of these results, we infer that the cross-linked sites identified by PAR-CLIP define the Hfq binding region on the *osmY* mRNA.

A second mRNA that was analyzed was *mdtK*, which was found to cross-link to Hfq 66 nts upstream of the initiation codon, and whose expression was found to be enhanced as adjudged by proteomics analysis. Consistent with the results of these analyses, we found that the expression of a *mdtK-lacZ* fusion was increased in a Δ*hfq* strain by 1.9-fold. We next mutated nine residues overlapping the cross-linking site and found that the regulation by Hfq was reduced significantly. These results suggest that the mutated residues largely define the region of the mRNA that is contacted by Hfq. However, since regulation was not completely abolished, some residues outside of the mutated region might also contribute to Hfq binding. Interestingly, we also noted that the mutant *mdtK-lacZ* fusion was expressed at lower levels in both strains, consistent with a positive effect of Hfq on *mdtK* expression.

A third RNA tested was *ygdQ*, which was found to be upregulated by RNA-Seq and was found to contain two cross-linking sites, one in the 5’ UTR and the other in the coding region. The *ygdQ-lacZ* fusion was found to be upregulated by RNA-Seq by nearly three-fold in the Δ*hfq* background, but mutations at the two cross-liking sites resulted in significant down-regulation of the Hfq-mediated response. In the additional seven fusions that were tested, mutation of between one to six nts around the cross-linking site reduced the extent of Hfq-mediated regulation in all but two examples (Fig. 5D).

Overall, the general conclusion is that the cross-linking sites identified by PAR-CLIP not only identify the specific sites of interaction with Hfq, but also to define the regions through which Hfq mediates its biological effects with high accuracy. To the best of our knowledge, a rigorous test of whether the contact sites identified via cross-linking methods pinpoint the sites responsible for RNA regulation has not been yet tested. These analysis, therefore, provides a strong validation for the use of PAR-CLIP to identify functionally relevant interaction sites. In summation, the use of PAR-CLIP appears to have high utility both to identify the cross-linking sites of RNA-BPs on a transcriptome-wide scale and the specific regions on the RNAs through which they mediate their regulatory effects.

## Discussion

PAR-CLIP was developed 10 years ago, but within a relatively short time, it has been implemented for use both in higher and lower eukaryotes (20). For reasons that are unclear, this technique has not been used in prokaryotes as yet. To determine whether PAR-CLIP can be used to characterize RNA binding by prokaryotic RNA-BPs, several experiments were performed. First, we found that the addition of 4SU to the growth medium results in the incorporation of this nucleoside into RNA. When cells were grown with 300 μM of 4SU in the medium, the fraction of 4SU containing residues incorporated into RNA was found to be 1.3% (Fig. 1), close to the 1.4-2.4% range of 4SU incorporation that has been reported for eukaryotic cells (8). Thus, the ability to apply PAR-CLIP in bacteria is not limited by the import of 4SU into cells or by discrimination against this base by prokaryotic RNA polymerase.

Next, to determine conditions suitable for implementing PAR-CLIP, we evaluated crosslinking efficiency in cells grown with different levels of 4SU. We found that the extent of crosslinking initially rose with increased 4SU concentration, but high 4SU concentrations caused a signal reduction (Fig. 2A). Concurrently, high 4SU concentrations were found to reduce the growth rate (Fig. 2E). The highest amount of crosslinking was observed with 500 μM 4SU in the medium, which therefore represents a concentration suitable for maximum crosslinking efficiency. Alternatively, if minimizing the effects of 4SU on cell growth is desired, PAR-CLIP can also be effectively performed on cells grown with 250 μM 4SU. We expect that a similar approaches will be useful to determine the optimum conditions for using PAR-CLIP in other prokaryotic organisms.

We next applied PAR-CLIP to a strain containing FLAG-tagged Hfq as well to a control untagged strain. A comparison between reads from the two strains showed that the ncRNA category exhibited the largest increase in T>C conversion ratios in the former strain (Fig 3A), affirming the central role of Hfq in interactions with sRNAs, with a corresponding decrease in mRNA reads. Furthermore, of the 22 sRNAs enriched in the Hfq libraries (Fig. 3B), only three (SroH, PsrD and SymR) are not established Hfq targets, suggesting a low level of false-positives regardless of whether these sRNAs are found to be Hfq targets in the future or not. Thus, the use of PAR-CLIP provides confidence that the application of this method to other proteins will yield a substantially reliable list of genuine RNA targets.

A specific advantage of PAR-CLIP over other CLIP methods is that it allows the identification of cross-linking sites at nucleotide resolution through the identification of diagnostic T>C conversion events. Using this criterion, we were able to identify one or more sites associated with an increased T>C conversion frequency in several sRNAs and mRNAs. Additionally, by RNA-Seq and proteomics analyses, we identified a large number of genes that were regulated by Hfq in midlog or stationary phase (Fig. 4; Supplementary Tables 1 and 3). For both the up-regulated and down-regulated genes, cross-linking sites in their mRNA UTRs or coding regions could be identified. Furthermore, for several of these mRNAs, no associated regulatory sRNAs have been identified. These observations suggest that Hfq may regulate gene expression through a variety of mechanisms that will require further investigation.

We further tested whether mutation of residues near the cross-linking site impacts regulation, and in almost every case we found that the extent of regulation by Hfq was significantly reduced, indicating the PAR-CLIP accurately identifies the *cis*-acting RNA sequences through which Hfq regulates the RNAs. Based on these results, we believe that the use of PAR-CLIP has the potential to yield detailed insights into the basis of mRNA regulation by RNA-BPs at a whole-genome level with single-nucleotide resolution.

## Methods

### Strains and plasmids

The wild-type strain used for this study is CJ2109, a derivative of MG1655 that contains a point mutation in *rph*, which restores the reading frame of this prematurely terminated gene (21). Plasmid pCJ1099 was constructed by amplifying chromosomal DNA using primers with the sequences 5’-AAAAAGCTTGTCATGGCAGGATTATTCATCG-3’ and 5’-TTGGATCCTACTTGTCATCGTCATCCTTGTAGTCGATGTCATGATCTTTATAATCACCGTCATGGTCTTTGTAGTCCTGCGCAGCGGCAGGTTTACGCGG-3’. The PCR product, which contains the RhlE gene and an appended 3X-FLAG-tag sequence, was digested with BamHI and HindIII and subcloned into the BamHI and HindIII-digested arabinose-inducible expression vector, pMPM-A4 (22). CJ2191 was made by transforming pCJ1099 into CJ2109. Strain CJ2192, was constructed by appending a 3X FLAG-tag sequence to the C-terminus of the chromosomal Hfq gene by using recombineering (23). Strain derivatives containing Δ*hfq*, Δ*lacZ* or Δ*pcnB* mutations were made by transduction of the respective alleles from the Keio strains collection into CJ2109 (24). *LacZ* fusion constructs were made by amplifying genes by PCR using *E. coli* DNA, digesting the amplified products with *Bgl*2 and *Eco*R1 or *Mfe*I, and subcloning the products into pLACZY1a digested with *Bam*H1 and *Eco*R1 (25). The sequence of the oligonucleotides used for PCR amplification are shown in Supplementary Table 5. Mutations into these plasmids were introduced using a Q5 site-directed mutagenesis kit (New England Biolabs).

### Analysis of 4SU incorporation into RNA

CJ2109 was grown in LB medium in the presence or absence of 300 μM 4SU and total RNA was isolated from mid-log cells using the hot-phenol method. 15 μg of each RNA sample was digested to nucleosides using snake venom phosphodiesterase and alkaline phosphatase, and analyzed by HPLC, as described (8).

### PAR-CLIP

25 ml of bacterial culture was grown to an OD_600_ of 0.5 in the absence or presence of 4SU added at early log-phase. For strains using CJ2191, cells were grown in LB medium supplemented with ampicillin, and arabinose was added to a final concentration of 0.2% at OD_600_ = 0.2. Cells were cooled rapidly by adding ice and pelleted in a centrifuge. The cell pellets were resuspended in 10 ml of ice-cold saline solution and crosslinked at 365 nm on ice for five min, or the indicated times, using a CL-1000 ultraviolet crosslinker (UVP). The cells were re-pelleted by centrifugation and stored at −80°C. For cells grown in RDM, cells were harvested at mid-log phase and crosslinked directly at 365 nm for five minutes (1.7 w/cm^2^). After cross-linking, the cells were transferred to cold 40 ml tubes containing 10 g of ice, pelleted by centrifugation and stored at −80°C. Thawed cells were resuspended in 200 μl NP‐T buffer (50 mM Sodium phosphate, 300 mM NaCl, 0.05% Tween, pH 8.0), sonicated and centrifuged in a microfuge at 4°C. The supernatant fraction was transferred to a new tube and an equal volume of NP-T buffer supplemented with 8 M urea was added. The supernatant was incubated at 65°C for five min, followed by the addition of additional NP-T buffer to yield a final urea concentration of 1 M. The lysate was added to pre-washed anti‐FLAG magnetic beads (15 μl of a 50% bead suspension; Sigma, cat # A2220) and the mixture was mixed with rotation for 1 h at 4°C. The beads were collected by centrifugation at 8000 × *g* for 30 sec, resuspended in 100 μl NP‐T buffer, and washed twice with 100 μl high‐salt buffer (50 mM NaH_2_PO_4_, 1 M NaCl, 0.05% Tween, pH 8.0) and twice with 100 μl NP‐T buffer. The beads were resuspended in 100 μl of NP‐T buffer containing 1 mM MgCl_2_ and 50 units of Benzonase (Sigma; cat # E1014) and incubated for 10 min at 37°C in a heat block. The beads were washed once with high‐salt NP‐T buffer, twice with CIP buffer (100 mM NaCl, 50 mM Tris–HCl pH 7.4, 10 mM MgCl_2_), resuspended in 100 μl CIP buffer with 10 units of calf intestinal alkaline phosphatase (NEB) and incubated for 30 min at 37°C in a heat block. The beads were washed once with high‐salt buffer, twice with PNK buffer (50 mM Tris–HCl, pH 7.4, 10 mM MgCl_2_, 0.1 mM spermidine), resuspended in 100 μl PNK buffer with 10 units of T4 polynucleotide kinase and 1 μCi of γ-^32^P-ATP, and incubated for 30 min at 37°C. The beads were washed thrice with NP-T buffer and resuspended in 20 μl Protein Loading buffer (62 mM Tris, pH 6.8, 2% SDS, 0.01% bromophenol blue, 10% glycerol) and incubated for 3 min at 95°C. The beads were pelleted by centrifugation, and the supernatant was loaded on a mini-Protean 4-15% SDS–polyacrylamide gel (Bio-Rad), along with pre-stained protein markers. The RNA–protein complexes were transferred to nitrocellulose membrane using a Mini Trans-blot electrophoretic transfer cell (BioRad). The protein marker bands were overlaid with radioactive ink, and the membrane was exposed overnight on a phosphorimager screen. The bands corresponding to labeled RNA– protein complexes were cut into small pieces and incubated at 37°C with 400 μl PK solution [50 mM Tris–HCl, pH 7.4, 75 mM NaCl, 6 mM EDTA, 1% SDS, 10 units of RNasin (Promega) and 400 μg of proteinase K (ThermoScientific)] for 30 min with shaking at 37°C. 100 μl of 9 M urea solution was added and the incubation was continued for an additional 30 min. 450 μl of the PK/urea solution was mixed with 450 μl phenol/chloroform/isoamyl alcohol (25:24:1) alcohol and centrifuged for 12 min (16,000 × *g*) at 4°C. The aqueous phase was precipitated by overnight incubation at −20°C after the addition of 3 volumes of ethanol, 1/10 volume of 3 M Sodium Acetate (pH 5.2) and 1 μl of GlycoBlue (Life Technologies). The precipitate was pelleted by centrifugation (30 min, 16,000 × *g*, 4°C), washed with 80% ethanol, centrifuged again (15 min, 16,000 × *g*, 4°C), dried and resuspended in 10 μl of sterile water.

### Sequence library preparation

High throughput sequencing libraries were prepared using RNA isolated from total RNA or PAR-CLIP samples using the NEBNext Multiplex Small RNA Library Prep Set for Illumina (Set 2; cat# E7580, New England Biolabs) according to the manufacturer’s instructions. To prepare RNA-Seq libraries, 1 μg of total RNA was digested with 0.01 units of Benzonase for 10 min at 37°C in 20 μl of NP-T buffer, followed by extraction with phenol/chloroform/isoamyl alcohol and ethanol precipitation. 100 ng of the digested RNA was used for sequence library preparation as described above. To prepare PAR-CLIP libraries, RNA isolated using the PAR-CLIP protocol was treated similarly.

### Bioinformatics Data Analysis

Raw sequencing data were analyzed by using the web-based Galaxy platform (13). Raw data was converted to Fastqsanger format and adaptors were removed using *Trim Galore!* The reads were aligned to the *E. coli* genome (Genbank: NC_000913.3) using *Bowtie 2*. For the analysis of RNA-seq data, uniquely aligned reads corresponding to individual RNAs were determined using *htseq-count*. For analysis of PAR-CLIP data, data from five biological replicates from strains CJ2109 and CJ2192 were pooled. T>C conversion frequencies at each genome coordinate were determined by using *Naïve Variant Caller* and *Variant Annotator.* Sites of Hfq-mediated cross-linking were assigned based on: (a) ≥ 5 reads from each library, (b) >0.1 conversion frequencies for data from strain CJ2192, (b) conversion frequencies in the data for strain CJ2192 ≥ twice the conversion frequencies in the data from strain CJ2109.

### Quantitative mass spectrometry

*E. coli* strains were grown in RDM to mid-log or stationary phase, sonicated and centrifuged. 100 μg of proteins from the supernatant fraction were dried under vacuum and processed for digestion using micro S-Traps™ (Protifi, Huntington, NY) according to the manufacturer’s instructions. Briefly, the dried protein was resuspended in 22 μl of 5% SDS, reduced using DTT at 55°C for 10 minutes, alkylated using methyl methanethiosulfonate (MMTS) at room temperature for 10 minutes and then spun down at 20K for 10 minutes. Subsequently, 2.5 μl of phosphoric acid was added to the sample, followed by 165 μl of a mixture of HPLC grade methanol and Protifi binding/wash buffer. Following loading onto the S-Trap™, samples were washed three times using centrifugation and trypsin was added in 50 mM TEAB at a 1:25 w/w ratio. The S-Trap™ column was incubated for 1 hour at 47°C. Following this incubation, 40 μl of 50 mM triethylammonium bicarbonate (TEAB) was added to the S-Trap™ and the peptides were eluted using centrifugation. Elution was repeated once. A third elution using 35 μl of 50% acetonitrile (ACN) was also performed and the eluted peptides were dried under vacuum. Peptides were reconstituted in 50 mM TEAB and their concentrations were determined using the Pierce™ quantitative fluorometric peptide assay (Thermo Fisher Scientific, Waltham, MA). Twelve micrograms of peptides were labelled with TMT labels (6-plex) according to the manufacturer’s instructions (Thermo Fisher Scientific, Waltham, MA) and pooled. The pooled plexed samples were dried under vacuum, resolubilized in 1% trifluoroacteic acid (TFA), desalted using 2 μg-capacity C18 ZipTips (Millipore, Billerica, MA), and then dried once again using vacuum. For mass spectrometry, dried TMT-labelled peptides were reconstituted in 5 μl of 0.1% TFA, vortexed briefly, and then sonicated for 15 minutes. The peptides were subsequently on-line eluted into a Fusion Tribrid mass spectrometer (Thermo Scientific, San Jose, CA) from an Acclaim PepMap™ RSLC C18 nano Viper analytical column (2 μM, 100Å, 75-μm ID × 50 cm, Thermo Scientific, San Jose, CA) using a gradient of 5-25% solvent B (80/20 ACN/water, 0.1% formic acid) in 180 min, followed by 25-44% solvent B in 60 min, 44-80% solvent B in 0.1 min, a 5 min hold of 80% solvent B, a return to 5% solvent B in 0.1 min, and finally a 20 min hold of solvent B. All flow rates were 250 nl/min delivered using a nEasy-LC1000 nano liquid chromatography system (Thermo Scientific, San Jose, CA). Solvent A consisted of water and 0.1% formic acid. Ions were created at 1.7 kV using the EASY-Spray™ ion source held at 50°C (Thermo Scientific, San Jose, CA). A synchronous precursor selection (SPS)-MS^3^ mass spectrometry method was used, as described (26), scanning between 380-2000 m/z at a resolution of 120,000 for MS1 in the Orbitrap mass analyzer, and performing collision-induced dissociation (CID) at top speed in the linear ion trap of peptide monoisotopic ions with charge 2-8, using a quadrupole isolation of 0.7 m/z and a CID energy of 35%. The top 10 MS^2^ ions in the ion trap between 400-1200 m/z were then chosen for higher energy collision dissociation (HCD) at 65% energy and detection in the Orbitrap at a resolution of 60,000 and an automatic gain control (AGC) target of 100,000 and an injection time of 120 msec (MS^3^). Quantitative analysis of the TMT experiments was performed simultaneously to protein identification using Proteome Discoverer 2.3 software (Thermo Scientific, San Jose, CA) using the raw data. The precursor and fragment ion mass tolerances were set to 10 ppm, 0.6 Da, respectively), enzyme was trypsin with a maximum of 2 missed cleavages and the Uniprot Escherichia Coli K12 proteome FASTA file was used in the SEQUEST searches. The impurity correction factors obtained from Thermo Fisher Scientific for each kit were included in the search and quantification. The following settings were used to search the data: oxidation, +15.995 Da (M); deamidation, +0.984 Da (N, Q); static modifications of TMT 6-plex +229.163 Da (N-terminus K); MMTS +45.988 (C). Only Unique+ Razor peptides were considered for quantification purposes. The Percolator feature of Proteome Discoverer 2.3 was used to set a false discovery rate (FDR) of 1%. Total Peptide Abundance normalization method was used to adjust for loading bias and the protein abundance based method was used to calculate the protein level ratios. Co-isolation threshold and synchronous precursor selection (SPS) mass matches threshold were set to 50 and 65, respectively. ANOVA was performed by individual proteins and the multiple testing correction as per the Benjamini-Hochberg procedure was applied to identify a “top tier” of significant proteins and a column for p_adj_ values was created.

### β-galactosidase assays

Plasmid constructs were transformed into Δ*lacZ* versions of strains CJ2109 or CJ2109 Δ*hfq*. In some instances, the strains contained an additional Δ*pcnB* mutation, since we showed that the Δ*pcnB* mutation reduces the toxicity associated with high levels of *lacZ* expression (27). Strains were grown in RDM (Teknova) and assayed in mid-log or saturated phase. Each measurement reported is based on four biological replicates.

## Acknowledgements

We thank Dr. Zhongwei Li (Florida Atlantic University) for assistance with HPLC analysis. We thank Thorsten Bischler and Sung-Huan yu (University of Würzburg) for providing a GFF file with 5’ UTR annotations. This work was supported by Grant GM114540 from the National Institutes of Health.

## Notes

### Competing Interest Statement

The authors have declared no competing interest.

